# Parvalbumin interneurons in the nucleus accumbens regulate impairment of risk avoidance in *DISC1* transgenic mice

**DOI:** 10.1101/2020.09.10.291450

**Authors:** Xinyi Zhou, Bifeng Wu, Qian Xiao, Wei He, Ying Zhou, Pengfei Wei, Xu Zhang, Yue Liu, Jie Wang, Weidong Li, Liping Wang, Jie Tu

## Abstract

One strong survival instinct in animals is to approach things that are of benefit and avoid risk. In humans, a large portion of mental disorders are accompanied by cognition-related impairments including the inability to recognize potential risks. One of the most important genes involved in risk behavior is disrupted-in-schizophrenia-1 (*DISC1*), and animal models where this gene has some dysfunction show cognitive impairments. However, whether *DISC1* mice models have an impairment in avoiding potential risks is still not fully understood. In the present study, we used DISC1-N terminal truncation (*DISC1-N^TM^*) mice to study cognitive abilities related to potential risks. We found that *DISC1-N^TM^* mice were impaired in risk avoidance on the elevated plus maze (EPM) test, and showed impairment in social preference in a three-chamber social interaction test. Staining for c-Fos following the EPM indicated that the nucleus accumbens (NAc) was associated with risk avoidance behavior in *DISC1-N^TM^* mice. Meanwhile, *in vivo* electrophysiological recordings showed that firing rates of fast spiking neurons (FS) in the NAc significantly decreased in *DISC1-N^TM^* mice following tamoxifen administration. In addition, theta band power was lower when mice shuttled from the safe (closed) arms to the risky (open) arms, an effect which disappeared after induction of the truncated *DISC1* gene. Furthermore, we found through *in vitro* patch clamp recording that the frequency of action potentials stimulated by current injection was lower in parvalbumin (PV) neurons in the NAc of *DISC1-N^TM^* mice than their wild-type littermates. Risk-avoidance impairments in *DISC1-N^TM^* mice were rescued using optogenetic tools that activated NAc^PV^ neurons. Finally, we inhibited activitiy of NAc^PV^ neurons in PV-Cre mice, which mimicked the risk-avoidance impairment found in the *DISC1-N^TM^* mice during tests on the elevated zero maze. Taken together, our findings confirmed a cognitive impairment in *DISC1-N^TM^* mice related to risk recognition and suggests that reduced excitability of NAc^PV^ neurons may be responsible.

## Introduction

Many patients with mental disorders are accompanied by varying levels of cognitive deficiency, which can influence psychosocial outcomes and raise the probability of having an accident [1]. The study of an extended Scottish family in which a balanced (1; 11) (q42.1;q14.3) chromosomal translocation was found to co-segregate with mental illness led the way to the identification of the DISC1 gene [2]. The *DISC1* gene is now perhaps the most well-known risk genes used to study the pathophysiology of major mental disorders, such as schizophrenia, bipolar disorder, and major depressive disorder (MDD) [3]. Many transgenic rodent models have been generated to manipulate the function of *DISC1* gene [4–6]. Many studies have shown that *DISC1* transgenic rodents show abnormal cognitive function, including object recognition memory deficits, working memory deficits, and social recognition deficits [7, 8]. These cognitive deficits are consistent with those found in human patients with mental disorders [9–11].

Another important cognitive ability is the evaluation of risk and reward, which is essential for survival, and as such, risk avoidance is a behavior conserved across species [12–14]. Although both genetic and functional studies provide evidence to support a critical role of *DISC1* in risk recognition, the underlying neuronal mechanism has not yet been discovered. We do know that abnormal development and impaired PV interneuron function have been found in mice with *DISC1* truncation [15, 16]. Previous work from our group has suggested that activity of NAc^PV^ neurons are required for anxiety-like behavior [17]. However, whether NAc^PV^ neuronal activity is also involved in risk cognition and avoidance has not been determined.

In the present study, we addressed these questions using a combination of *DISC1* truncation transgenic mice, opto/chemogenetic manipulations, *in vivo* recordings and behavioral analyses to examine whether the cognitive deficiency phenotype in *DISC1-N^TM^* mice encompasses impairment of risk recognition and also to investigate the underlying neuronal mechanisms.

## Materials and Methods

### Animals

We used the following mouse lines: PV-Cre mice (B6; 129P2-Pvalbtm1 (cre) Arbr/J, Jackson Laboratory, stock No.008069), VGAT-ChR2-EYFP (B6.Cg-Tg (Slc32a1-COP4*H134R/EYFP) 8Gfng/J, Jackson Laboratory, stock No. 014548), inducible *DISC1-N* terminal fragment transgenic mice (*DISC1-N^TM^*, from Prof. Weidong Li, Shanghai Jiao Tong University) and C57BL/6J mice (Guangdong Medical Laboratory Animal Center, Guangzhou, China). Mice were given free access to food pellets and water and maintained on a 12h/12h light/dark cycle (lights on at 8:00 a.m.). All experiments were approved by the Shenzhen Institutes of Advanced Technology, Chinese Academy of Sciences Research Committee, and all experimental procedures involving animals were carried out in strict accordance with the Research Committee’s animal use guidelines. Surgeries were performed under full anesthesia and every effort was made to minimize animal suffering.

### Behavior tests

All behavioral tests were performed blind to mice genotype. Groups of mice were age-matched (8-16 weeks) and, prior to behavioral assays, were handled for 5 min per day for three days to reduce stress introduced by experimenter contact. *DISC1-N^TM^* mice were given tamoxifen (Sigma-Aldrich, #T5648) (intraperitoneal injection, 5X10^−5^g/g) two hours before behavioral testing and finished testing within 6 hours. Behavioral tests using CNO (MedChemExpress, #HY-17366) were conducted in a 60-min window that began 30 min after CNO administration (intraperitoneal injection, 1mg/kg). All behavioral tests were recorded by a video camera directly above and videos were analyzed using ANY-maze (Stoelting, U.S.A).

#### 1) Elevated plus maze test (EPM)

A plastic elevated plus maze consisting of a central platform (5×5 cm) with two white open arms (25×5 cm) and two white closed arms (25×5 cm, with 17-cm-high surrounding walls) extending from it in a plus shape was used. The maze was elevated 58 cm above the floor. Mice were placed in the central area of the EPM with their heads facing an open arm and were allowed to freely explore for 5 min. The number of entries and the amount of time spent in each arm type were recorded.

#### 2) Elevated zero maze test

A plastic elevated zero maze consisting of an elevated annular platform (external diameter: 50 cm, width: 5 cm) with two opposite, enclosed quadrants (height: 15 cm) and two open quadrants was used. The maze was elevated 80 cm above the floor. Mice were placed on the platform with their heads facing one closed arm and behavior was recorded for 5 min. Similar to the EPM, the number of entries and the amount of time spent in each arm type were recorded.

#### 3) Three-chamber social interaction test

A three-chambered apparatus (60×40×25 cm) with a central chamber (20-cm wide) and two side chambers (each 20-cm wide) was used. An empty housing cage was put inside each side chamber and the test mouse was placed in the central chamber and allowed to freely explore the three chambers for a 5 min habituation period. Then, in a second phase, the mouse was gently put back to the central chamber and the two side-chamber entrances were blocked and then a ‘stranger’ mouse was placed in one side chamber. Both entrances were then opened to allow the test mouse to explore the new environment freely for 10 min. The time spent in each chamber was recorded. All stranger mice were of the same age and gender.

### Stereotactic virus injection and optogenetic manipulation

Adeno-associated viruses (AAVs) carrying Cre-inducible transgenes (AAV-DIO-ChR2-mCherry, titers 3×10^12^ particles per ml, AAV-DIO-hM4Di-mCherry, titers 3×10^12^ particles per ml, AAV-DIO-mCherry, titers 3×10^12^ particles per ml) were packaged in our laboratory. Mice were deeply anesthetized with 1% sodium pentobarbital (Sigma-Aldrich, #P3761, 10 ml/kg body weight, i.p.) and placed in a stereotaxic instrument (RWD Life Science Inc., Shenzhen, China) and head fixed. A microinjector pump (UMP3/Micro4, USA) was used through a microliter syringe (10 μl, Hamilton) to inject virus to the target region at a speed of 80 nl/min. The needle was left in place for 10 min after finishing injection to avoid reflux of the viral solution. A volume of 200 nl of AAV-DIO-ChR2-mCherry was unilateral injected at NAc (AP +1.12 mm, ML +1.50 mm, DV −4.85 mm) for optogenetic and electrophysiological experiments. Similarly, 200 nl of AAV-DIO-hM4Di-mCherry was bilateral injected at NAc (AP +1.12 mm, ML ±1.50 mm, DV −4.85 mm) for designer receptors exclusively activated by a designer drug (DREADD) experiments. For optogenetic experiments, 2 weeks after virus injection, mice were implanted with a 200 μm unilateral fiber optic cannula (AP +1.12 mm, ML +1.50 mm, DV −4.65 mm) secured to the skull with denture base material (SND, China) and dental base acrylic resin powder (Feiying, China). The mice were then given 1-week recovery before behavior experiments began. A control (mCherry) group underwent the same procedure and received the same intensity of laser stimulation.

### Immunohistochemistry

Mice received an overdose of 1% sodium pentobarbital (15 ml/kg body weight, i.p.) and were transcardially perfused with phosphate-buffered saline (PBS), followed by 4% paraformaldehyde (PFA; Aladdin, #C104188) in PBS. Brains were removed and submerged in 4% PFA at 4 °C overnight to post-fix, and then transferred to 20% sucrose for one day and then 30% sucrose for 2 days. Coronal brains sections (30 μm) were obtained on a cryostat microtome (Lecia CM1950, Germany). Brain sections were washed with PBS (3 min, room temperature) 3 times to wash out OCT. Then, brain sections were put into blocking solution (0.3% Triton X-100 and 10% normal goat serum, NGS in PBS, 1 h at room temperature). Brain sections were then incubated in primary antiserum (rabbit anti-c-Fos, 1:300, Cell Signaling; rabbit anti-PV, 1:300, Abcam; mouse anti-HA-tag, 1:300, proteintech) diluted in PBS with 3% NGS and 0.1% Triton X-100 overnight. The following day, the sections were incubated in secondary antibodies at room temperature for 1 h. The secondary antibodies used were Alexa Fluor^®^ 488 or 594-conjugated goat anti-rabbit or anti-mouse IgG antibodies (1:300, Invitrogen, CA, USA) at room temperature for 1 h. Then brain sections were mounted and covered slipped with anti-fade reagent with DAPI (ProLong Gold Antifade Reagent with DAPI, life technologies). Brain sections were then photographed and analyzed with an Olympus slide scanner VS120-S6-W or Leica TCS SP5 laser scanning confocal microscope. Imagines were taken and c-Fos staining was manually counted by two individual experimenters blind to the experiment groups. The Mouse Brain in Stereotaxic Coordinates was used to locate brain areas. For c-Fos staining following behavioral tests, mice were sacrificed 1.5 hr post EPM stimulus and brains then subjected to c-Fos staining. For c-Fos staining without behavioral tests, mice were sacrificed directly after moving from homecage.

### Patch-clamp electrophysiology

For patch clamp recording, all drugs used were from Sigma-Aldrich unless indicated otherwise. Coronal slices (300 μm) containing NAc shell (bregma 1.7 to 0.6 mm) were prepared from mice using standard procedures. Brains were quickly removed and chilled in ice-cold modified artificial cerebrospinal fluid (ACSF) containing (in mM): 110 Choline Chloride, 2.5 KCl, 1.3 NaH_2_PO_4_, 25 NaHCO_3_, 1.3 Na-Ascorbate, 0.6 Na-Pyruvate, 10 Glucose, 2 CaCl_2_, 1.3 MgCl_2_. Then the NAc slices were cut in ice-cold modified ACSF using a Leica vibroslicer (VT-1200S). Slices were allowed to recover for 30 min in a storage chamber containing regular ACSF at 32~34 °C (in mM): 125 NaCl, 2.5 KCl, 1.3 NaH_2_PO_4_, 25NaHCO_3_, 1.3 Na-Ascorbate, 0.6 Na-Pyruvate, 10 Glucose, 2 CaCl_2_, 1.3 MgCl_2_ (pH 7.3~7.4 when saturated with 95% O_2_/5% CO_2_), and thereafter kept at room temperature, until placed in the recording chamber. The osmolarity of all the solutions was 300~320 mOsm/kg. For all electrophysiological experiments, slices were viewed using infrared optics under an upright microscope (Eclipse FN1, Nikon Instruments). The recording chamber was continuously perfused with oxygenated ACSF (2 ml/min) at room temperature. Pipettes were pulled by a micropipette puller (Sutter P-2000 Micropipette Puller) with a resistance of 5-10 MΩ. Recordings were made with electrodes filled with intracellular solution (in mM): 130 potassium gluconate, 1 EGTA, 10 NaCl, 10 HEPES, 2 MgCl_2_, 0.133 CaCl_2_, 3.5 Mg-ATP, 1 Na-GTP. Action potential firing frequency was analyzed in current-clamp mode in response to a 2 s depolarizing current step. All recordings were conducted with a MultiClamp700B amplifier (Molecular Devices). Currents were low-pass filtered at 2 kHz and digitized at 20 kHz using an Axon Digidata 1440A data acquisition system and pClamp 10 software (both from Molecular Devices). Series resistance (Rs) was10~30 MΩ and regularly monitored throughout the recordings. Data were discarded if Rs changed by >30% over the course of data acquisition.

### In vivo single-unit and local field potential (LFP) recordings in the NAc shell of *DISC1-N^TM^* mice

Microdrive tetrodes were constructed using platinum-iridium wire (12μm, California Fine Wire Co.). Eight tetrodes were arranged into a 4×2 array and attached to a microdrive such that the tetrodes could be driven towards a desired location. The impedance of the electrodes was reduced to around 0.5 MΩ by platinum electroplating.

*DISC1-N^TM^* mice (aged 8-12 weeks) were anesthetized with sodium pentobarbital (70 mg/kg i.p.). Body temperature was maintained at 37 °C with a heating pad placed under the mouse. The head of the mouse was secured and leveled in both the anterior-posterior and the middle-lateral axis on a stereotaxic apparatus. A microdrive electrode was attached to a micromanipulator and the electrodes were inserted gradually into the medial part of the NAc. The tetrode tips were ultimately aimed at the following coordinates: AP1.12 mm, ML1.50 mm, and DV-4.65 mm but were left 400 μm above the required depth during the surgery. The microdrive and connector were secured on the skull using dental cement. Mice were allowed to recover for at least one week after implantation.

*In vivo* extracellular signals were monitored and acquired using a 64-channel Multichannel Acquisition Processor (Plexon Inc, Dallas, TX) with a headstage amplifier (Plexon Inc, HST/32V-G20). Wideband signals were recorded at 40 kHz. Local field potential (LFP) signals were filtered at 0-200 Hz by a low-pass Bessel filter and downsampled to 1 kHz.

Prior to in vivo recording sessions, mice were habituated to the recording set-up with two daily 20-min sessions in their home cages where their microdrives were tethered to the recording cables in order to habituate to the plugging in/out process and to the overhead cable movements. Electrodes were gradually advanced to the desired depth and in vivo recordings were acquired when well-isolated single units were detected and were stable for at least 2 hours. Mice were allowed to freely explore the EPM for 5 min before and 6 hours after tamoxifen injection. Behavioral activity was recorded by a camera directly above and synchronized with neural data acquisition. After in vivo recording, recording sites were marked with electrolytic lesions and animals were deeply anesthetized with 10% chloral hydrate (0.4mg/kg) and transcardially perfused with PBS, followed by 4% paraformaldehyde (PFA) (wt/vol). Brains were dissected and post fixed at 4 °C in 4% PFA overnight. Frozen sections of the whole area were cut into 40-μm thick slices and the recording locations were reconstructed.

### Single-unit spike sorting and analysis

Mouse locomotion was analyzed and moving trajectories were reconstructed using an automatic video tracking system (AnyMaze). An offline sorter (Plexon) was used for spike detecting and spike sorting. Continuous wideband signals were high-pass filtered (300 Hz) with a Bessel filter, then a threshold (−4.5 s.d.) was applied for spike detection. A spike waveform window was set as 1400 μs including 300 μs before threshold crossing. Single units were sorted based on waveform features in three-dimensional principal component space. To quantify recording stability before and after tamoxifen injection, we calculated the correlation coefficient between the average waveforms of single units in the pre-injection session and post-injection session. Only cells exhibiting r values > 0.90 between the two sessions were recognized as the same cell and reserved for further analysis [18]. Single units in the NAc were classified into fast-spiking neurons and non-fast-spiking neurons utilizing an unsupervised clustering method using peak-to-peak width and averaged firing rate [19–21]. Burst number was calculated for each single unit where a burst was defined as composing at least 5 spikes with inter-spike interval (ISI) of 6 ms [22]. To compare firing rate changes before and after tamoxifen injection, the firing rate of each single unit was calculated in five successive bins (10 s per bin). Firing rate was recognized as undergoing a significant change when the P-value of the rank sum test was less than 0.05. The multitaper method [23] in the Chrounux analysis package (http://chronux.org/) was used for spectral analysis to estimate spectral power, time frequency and coherence analysis. The value was calculated using a 1 s window with 3 time-bandwidth product (NW) and 5 tapers. The coherence value in the theta band (4-8 Hz) that exceeded the 95% confidence level was used for analysis. The coherence value was normalized by dividing by the max value in the theta band. Statistical analysis of coherence was conducted on the original values.

## Results

### *DISC1-N^TM^* mice have impaired risk-avoidance behavior

The *DISC1* transgenic mouse strain used was an N-terminal fragment isoform under a *Camk2a* promoter (Figure 1A). To check the expression of the inserted gene, the HA-tag was co-stained with DNA-specific fluorescent Hoechst 33258 in three typical brain areas: the cortex, the hippocampus and the NAc (Figure 1B). This revealed that the *DISC1-N* truncation was widely expressed in the transgenic mice.

**Figure 1.**
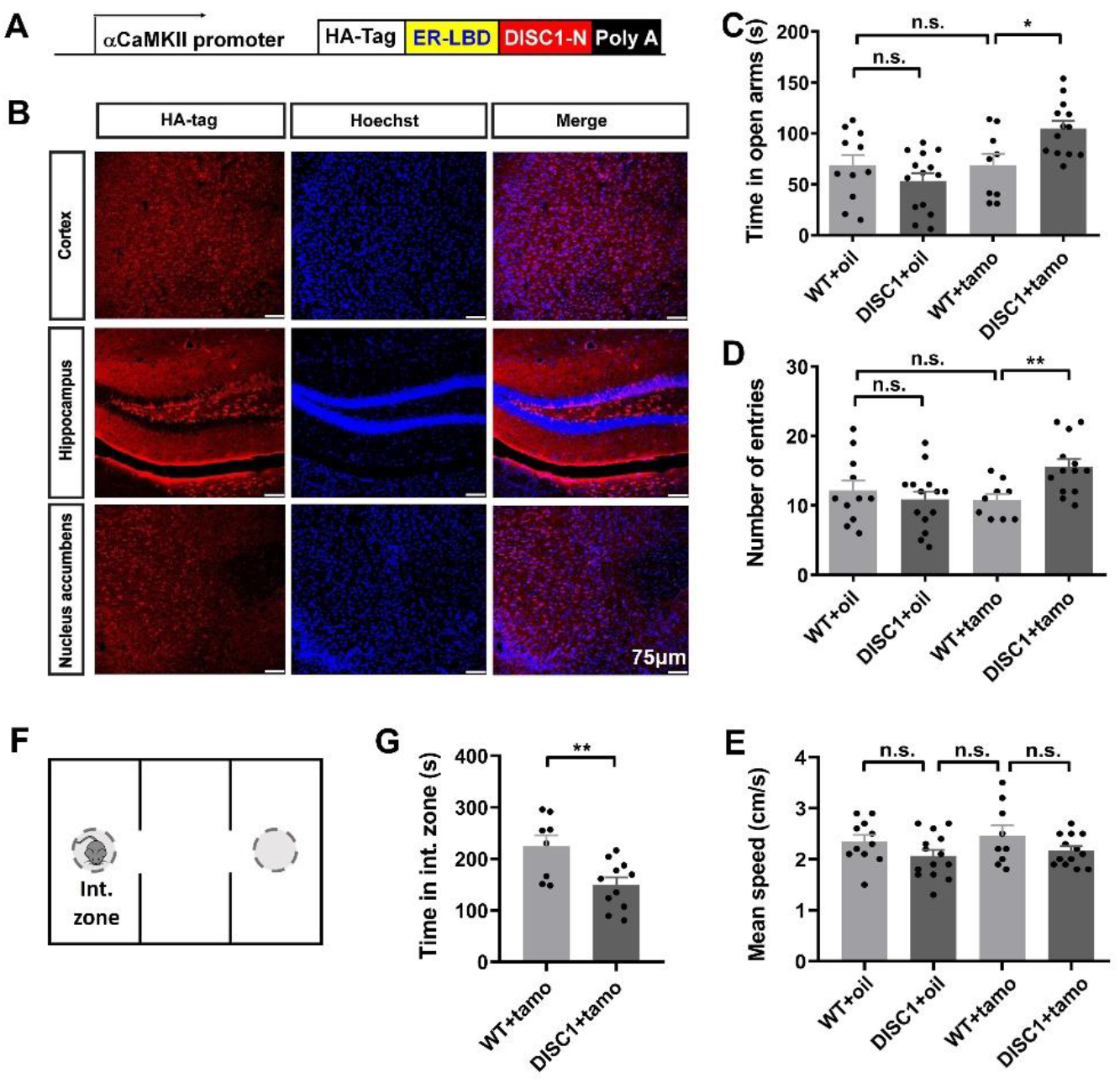
*DISC1-N^TM^* mice are impaired in risk avoidance. A. Schematic showing the inserted expressed sequence in *DISC1-N^TM^* mice. B. Staining for HA-tag (red) and hoechst 33258 (blue) in the cortex, hippocampus and NAc of *DISC1-N^TM^* mice. C-E. Time spent in the open arms (C), number of entries to the open arms (D) and mean speed (E) during the EPM test (unpaired t-test, \**P* =0.0105, \*\**P* =0.0058; n=11, 14, 9 and 13 from left to right). F. Schematic for three-chamber social interaction task. G. Time spent in the interaction zone during the three-chamber social interaction task (unpaired t-test, \*\**P* =0.0077; n=8 and 11 from left to right).

We next asked if expression of the N-terminal fragment isoform of the *DISC1* gene affected behavior related to risk recognition in the elevated plus maze (EPM). After tamoxifen administration (i.p.), *DISC1-N^TM^* mice spent significantly more time in the open arms and made more entries to the open arms compared to control groups including wild type with tamoxifen mice (WT+tamo), wild type with vehicle mice (WT+oil) and DISC1 mice with vehicle (DISC1+oil) (Figure 1C, D). There were no significant differences between the four groups in mean speed (Figure 1E). Furthermore, there was no difference between the DISC1+tamo and WT+tamo groups in velocity in the open or closed arms, total distance traveled, or time spent in the center (P > 0.05; Table 1). Therefore, the difference in residence time and number of entries to the open arms were unlikely to have be confounded by changes in locomotion or by intraperitoneal administration. These results suggest that the expression of the N-terminal fragment isoform of the *DISC1* gene influenced cognition of risk in the EPM test.

**Table 1.**
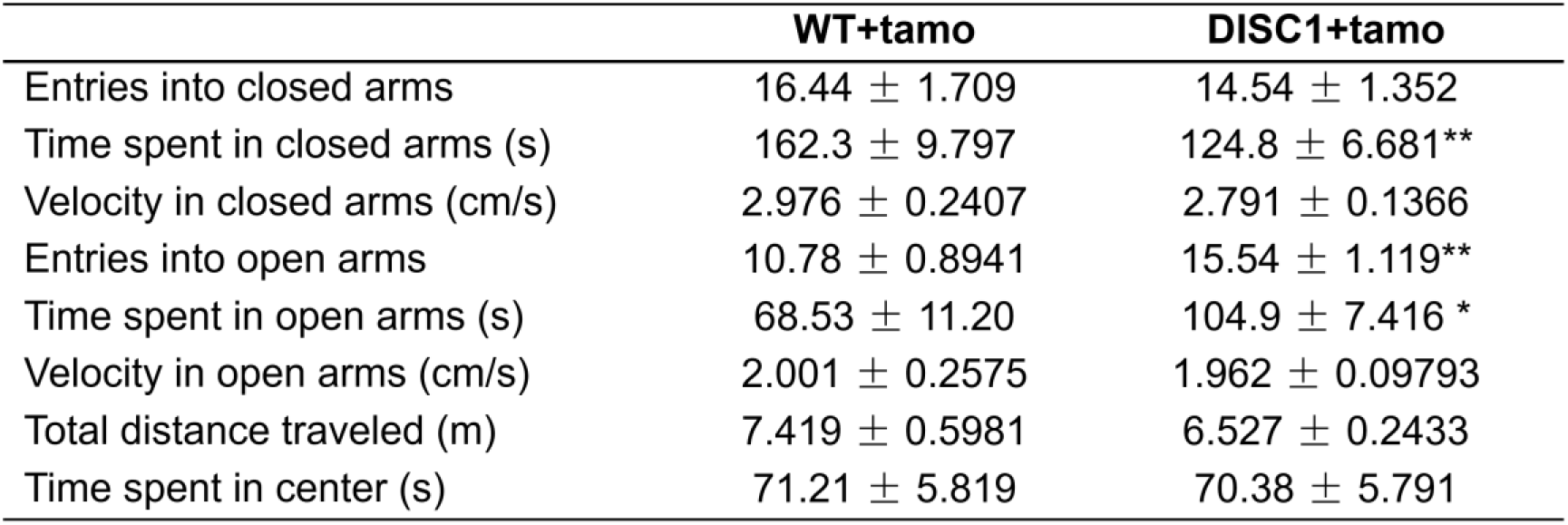
Results of elevated plus-maze test. Values are presented as mean ± SEM (\**P* =0.0105, \*\**P* =0.0037 and 0.0058 from top to bottom).

Because some *DISC1* mutant mice also show cognitive defects in social-related behavior [24], we used a three-chamber social interaction test to determine sociability in *DISC1-N^TM^* mice. Mice normally prefer to spend more time with other mice [25]; however, following tamoxifen injections, *DISC1-N^TM^* mice spent less time with a stranger mouse than did the WT+tamo control group (Figure 1G). Low interaction time with the stranger mouse indicates that *DISC1-N^TM^* mice had a social preference impairment.

### The NAc may play a role in the abnormal risk-avoidance behavior seen in *DISC1-N^TM^* mice, as illustrated by c-Fos staining

After confirming cognitive impairment in risk avoidance in *DISC1-N^TM^* mice, we investigated which brain area is involved in abnormal risk-avoidance behavior in *DISC1-N^TM^* mice. We performed c-Fos staining following an EPM test in eight cognition-related or emotion-related brain areas: the medial prefrontal cortex (mPFC), NAc, bed nucleus of the stria terminalis (BNST), hippocampus, lateral hypothalamic (LH), basolateral amygdala (BLA), paraventricular nucleus of the thalamus (PVT) and the ventral tegmental area (VTA). We found that c-Fos expression in the NAc, BLA, PVT and VTA was significantly elevated in *DISC1-N^TM^* mice compared to WT mice (Figure 2A). Our previous work showed that neurons in the NAc can control behavioral states and regulate mouse performance on the EPM [17]. Moreover, the NAc is considered to be an interface between cognition and emotion [26]. Thus, we hypothesized that the NAc may play a role in abnormal risk-avoidance behavior in *DISC1-N^TM^* mice.

**Figure 2.**
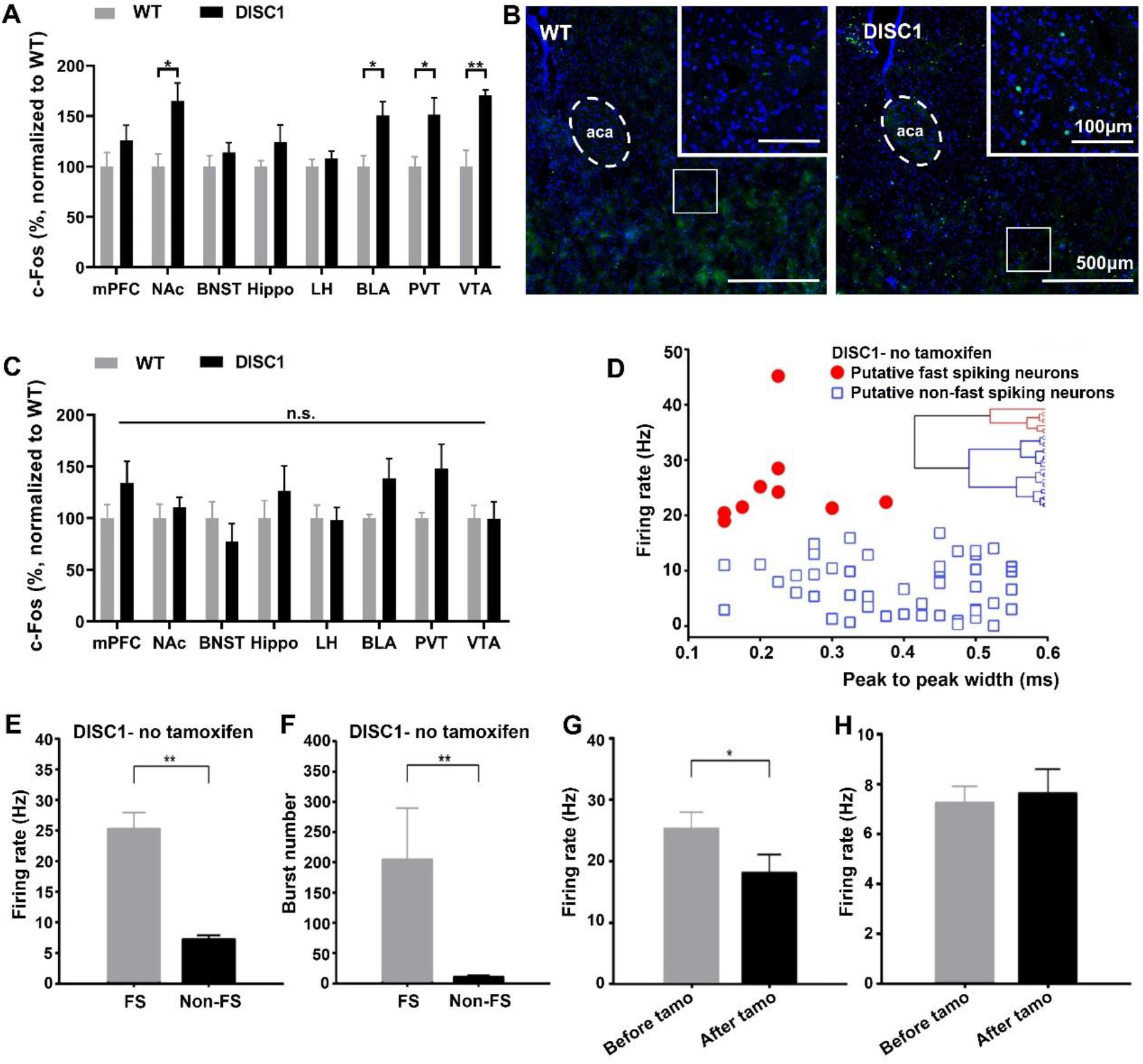
*In vivo* electrophysiological recordings in the NAc shell of *DISC1-N^TM^* with or without tamoxifen induction. A. Comparison of c-Fos expression in WT and *DISC1-N^TM^* mice following EPM test (unpaired t-test, \**P* =0.0138, 0.0152 and 0.0247 from left to right, **P= 0.0019; n=3 for each group). B. Representative staining for c-Fos expression after EPM task in the NAc. C. Comparison of c-Fos expression in WT and *DISC1-N^TM^* homecage controls (n= 3, each group). D. Scatter plot showing the firing rate and peak-to-peak width for 61 units from 21 *DISC1-N^TM^* mice. E, F. Firing rate (E) and burst numbers (F) of putative fast spiking (FS) neurons and putative non-fast spiking neurons (Non-FS) (unpaired t-test, ** P= 0.001). G. Firing rates of FS before and after tamoxifen injection (paired t-test, n = 9, * P < 0.05). H. Firing rates of Non-FS neurons before and after tamoxifen injection (paired t-test, n = 52, P = 0.963). Data are presented as mean ± SEM.

To control for differences due to mouse strain, we also performed c-Fos staining on the brains of homecage-only mice and found that the differences between WT mice and *DISC1-N^TM^* mice diminished (Figure 2C). This strongly suggests that the elevated c-Fos expression in the NAc was due to the effects of the EPM. In addition, c-Fos expression was lower in homecage mice than mice in the EPM experiment (both in *DISC1-N^TM^* mice and WT mice, data not shown).

### *In vivo* electrophysiological recordings show reduced firing rate in FS neurons after tamoxifen injection

To further investigate which cell-types in the NAc may participate in risk-avoidance impairment in *DISC1-N^TM^* mice, we conducted *in vivo* single-cell recording in the NAc before and after *DISC1-N^TM^* mice were given tamoxifen. Distinct sub-types of NAc neurons were classified based on their major electrophysiological properties [19, 27]. Of 61 recorded NAc units in the *DISC1-N^TM^* mice, neurons were classified as putative FS neurons and putative non-fast spiking (Non-FS) neurons according to firing rate and peak-to-peak width (Fig. 2D). Nine units were identified as putative FS neurons and 52 units were identified as Non-FS neurons. The firing rate and burst numbers of the putative FS neurons and the Non-FS neurons were significantly different (Fig.2E, F), which indicates that they were indeed two different classes of neuron as described previously [19]. The average firing rates of the FS neurons decreased and there was a significant difference before and after tamoxifen injection (Fig. 2G). For the Non-FS neurons, firing rates were not different before and after tamoxifen injection (Fig. 2H). In summary, *in vivo* single-cell recordings show that the firing rates of NAc-FS neurons show decreased after tamoxifen injection in *DISC1-N^TM^* mice.

### NAc theta power undergoes no discriminative changes correlated with the location on the EPM in *DISC1-N^TM^* mice

Theta oscillation plays a role in cognitive behavior [28] and is correlated with cognitive impairments [29, 30]. We investigated local NAc LFP in *DISC1-N^TM^* mice to see how theta oscillation changes during an EPM test. Before tamoxifen injection, theta (4-8 Hz) power was significantly lower on the open arms than that on the closed arms (Fig. 3A), which is consistent with our previously reported findings in wild-type mice [17]. However, after tamoxifen injection, LFP power in the theta band was not predictive of whether the *DISC1-N^TM^* mice were on the open arms or closed arms (Fig. 3B). A representative time-frequency map (Fig. 3C) also confirmed that theta power in *DISC1-N^TM^* mice was significantly reduced when traversing from the closed arms to open arms before tamoxifen injection, but after tamoxifen induction, theta power did not show a reduction between arms.

**Fig 3.**
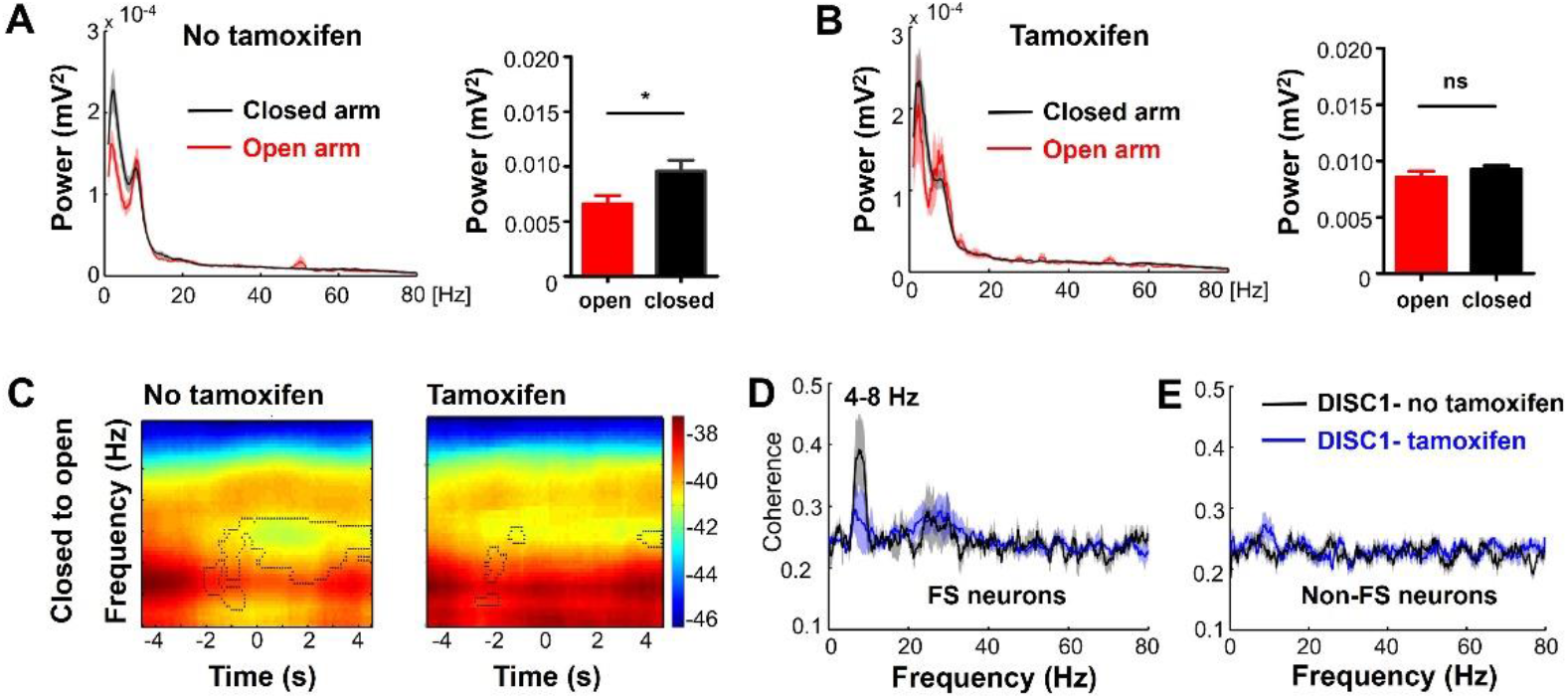
Accumbal LFP recordings in *DISC1-N^TM^* mice with or without tamoxifen induction. A. Changes in theta power during periods of traversing between different EPM arms types before tamoxifen induction (Paired *t*-test, n of experiments = 17, *t* = 2.192 with 16 degrees of freedom, \**P* = 0.044). B. No change in theta power during periods of traversing between different EPM arms types after tamoxifen induction (Paired *t*-test, n of experiments = 17, *t* = 0.119 with 16 degrees of freedom, *P* = 0.907). C. Example spectrograms showing the changes in LFP during animals traversing from the closed to open arms before and after tamoxifen injection in *DISC1-N^TM^* mice. D. Difference in spike and LFP coherence of PV-FS neurons in *DISC1-N^TM^* mice before and after tamoxifen injection (Wilcoxon rank-sum test, n = 4, W= −10, Z-Statistic = −1.826, P =0.0125). E. Difference in spike and LFP coherence in Non-FS neurons in *DISC1-N^TM^* mice before and after tamoxifen injection (Paired t-test, n = 13, t = 0.713 with 12 degrees of freedom, P = 0.488).

We then investigated coherence between spikes and LFP power. Coherence between FS neuron spikes and theta power was significantly lower in the *DISC1-N^TM^* mice (Fig. 3D). Conversely, for Non-FS neurons, there were no significant changes in coherence between spikes and theta power before or after tamoxifen injection (Fig. 3E). These data indicate that activity of NAc-FS neurons is strongly correlated with decrease of local theta power during exploration of the risky open arm. Taken together, these *in vivo* results indicate that theta power in the NAc was lower in the risky open arms and was correlated with the firing rate of NAc-FS neurons, while these phenomena were abolished by the induced expression of the N-terminal fragment isoform of *DISC1* gene. The firing rate of NAc-FS neurons was also lower after tamoxifen injection in the *DISC1-N^TM^* mice (Fig. 2G), suggesting that NAc-FS neurons may be involved in abnormal risk avoidance behavior in *DISC1-N^TM^* mice. One FS neuron type, PV interneurons, mediate theta oscillation in cortical networks [31]. What’s more, our previous work has shown that PV interneurons in the NAc regulate the performance of chronically stressed mice in the EPM [17]. Thus, we propose that PV interneurons in the NAc are response for the abnormal risk avoidance behavior of *DISC1-N^TM^* mice.

### PV neurons in the NAc show less excitability in *DISC1-N^TM^* mice

To further explore PV function in *DISC1-N^TM^* mice, we investigated the electrophysiological characteristics of NAc^PV^ neurons using the whole-cell patch clamp recordings. We used *PV-Cre* and *DISC1-N* truncation double transgenic mice and stereotaxically injected AAV-DIO-ChR2-mCherry into the NAc area. A PV-Cre mice control group was stereotaxically injected with AAV-DIO-ChR2-mCherry into the NAc area. After waiting 3-4 weeks for the virus to fully express, we prepared acute brain slices and recorded PV neurons under the current clamp model. Fluorescent mCherry, carried by the virus, enabled identification of PV neurons under the microscope (Figure 4B).

**Figure 4.**
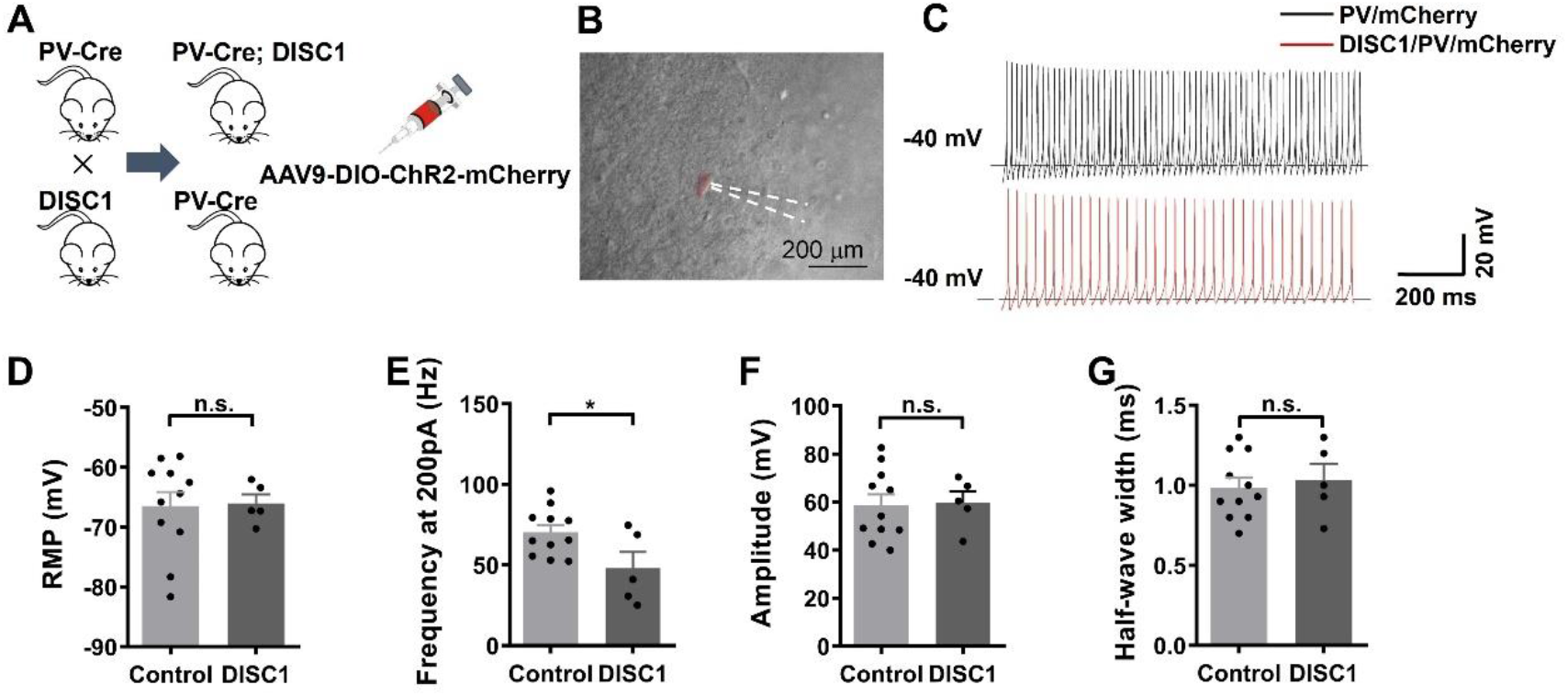
Recordings in acute brain slices from PV-Cre/*DISC1-N*^TM^ and PV-Cre mice. A. Schematic showing transgenic mice used. B. Representative photograph showing patch recording on a PV neuron. C. Representative plot showing action potentials stimulated by 200 pA current. D. Resting membrane potential of PV neurons (n=11 left, n=5 right). E-G. Frequency (E), amplitude (F) and half-wave width (G) of action potentials when giving 200 pA current stimulation (\**P* =0.0308, n=11 left, n=5 right).

We gave different step-current stimulation to the PV neurons using 100 pA, 150 pA and 200 pA. Only 200 pA current stimulation was able to induce stable action potentials (AP) in all PV neurons. So, we compared firing rates of the control and *DISC1-N^TM^* groups under 200 pA current stimulation and found that PV neurons in *DISC1-N^TM^* mice had significantly lower firing rate than PV-Cre control group (Figure 4C, E). This result indicates that NAc^PV^ neurons were less excitable than in *DISC1-N^TM^* mice than in control mice and that they may be responsible for risk-avoidance behavior impairment in *DISC1-N^TM^* mice. Resting membrane potential (Figure 4D), amplitude and half-wave width of APs evoked by a 200 pA current (Figure 4F, G) were not different between NAc^PV^ neurons of *DISC1-N^TM^* and PV-Cre control mice.

### Optogenetic activation of PV neurons in the NAc rescues risk avoidance impairment in *DISC1-N^TM^* mice

The results above indicate that NAc^PV^ neurons are dysfunctional in *DISC1-N^TM^* mice. Next, we tested whether modulation of PV neurons in the NAc could rescue the risk-avoidance impairments in *DISC1-N^TM^* mice. We selectively activated NAc^PV^ neurons by delivering blue light (5 ms per pulse, 60 Hz) in PV-Cre and *DISC1-N* truncation double transgenic mice unilaterally infected in the NAc with AAV-DIO-ChR2-mCherry (Figure 5A, B). PV-Cre mice unilaterally infected in the NAc with AAV-DIO-mCherry or AAV-DIO-ChR2-mCherry were also used as control groups (Figure 5A). The function of ChR2-expressing PV neurons had been checked by whole-cell patch clamp in brain slice containing the NAc area from PV-Cre mice (Figure 5C, D). During blue light stimulation, *DISC1-N^TM^* mice not only spent significantly less time in the open arms but also had lower entry numbers to the open arms compared with the light-off phase (Figure 5E, F). Furthermore, in the light-on phase, ChR2-expressing *DISC1-N^TM^* mice were not different from the non-ChR2-expressing PV-Cre control group in time spent in, and number of entries to the open arms (Figure 5E, F). These results suggest that PV neurons in the NAc can rescue abnormal risk-avoidance behavior in *DISC1-N^TM^* mice in the EPM task. Regarding the ChR2-expressing PV-Cre control group, when given blue light stimulation, time spent in and the number of entries to the open arms did not differ from the no-light state (Figure 5E, F). Activation of NAc^PV^ neurons in mice with normal *DISC1* gene expression did not significantly influence behavior. This indicates that in normal mice, PV neurons may play a role in maintaining an appropriate risk-avoidance state.

**Figure 5.**
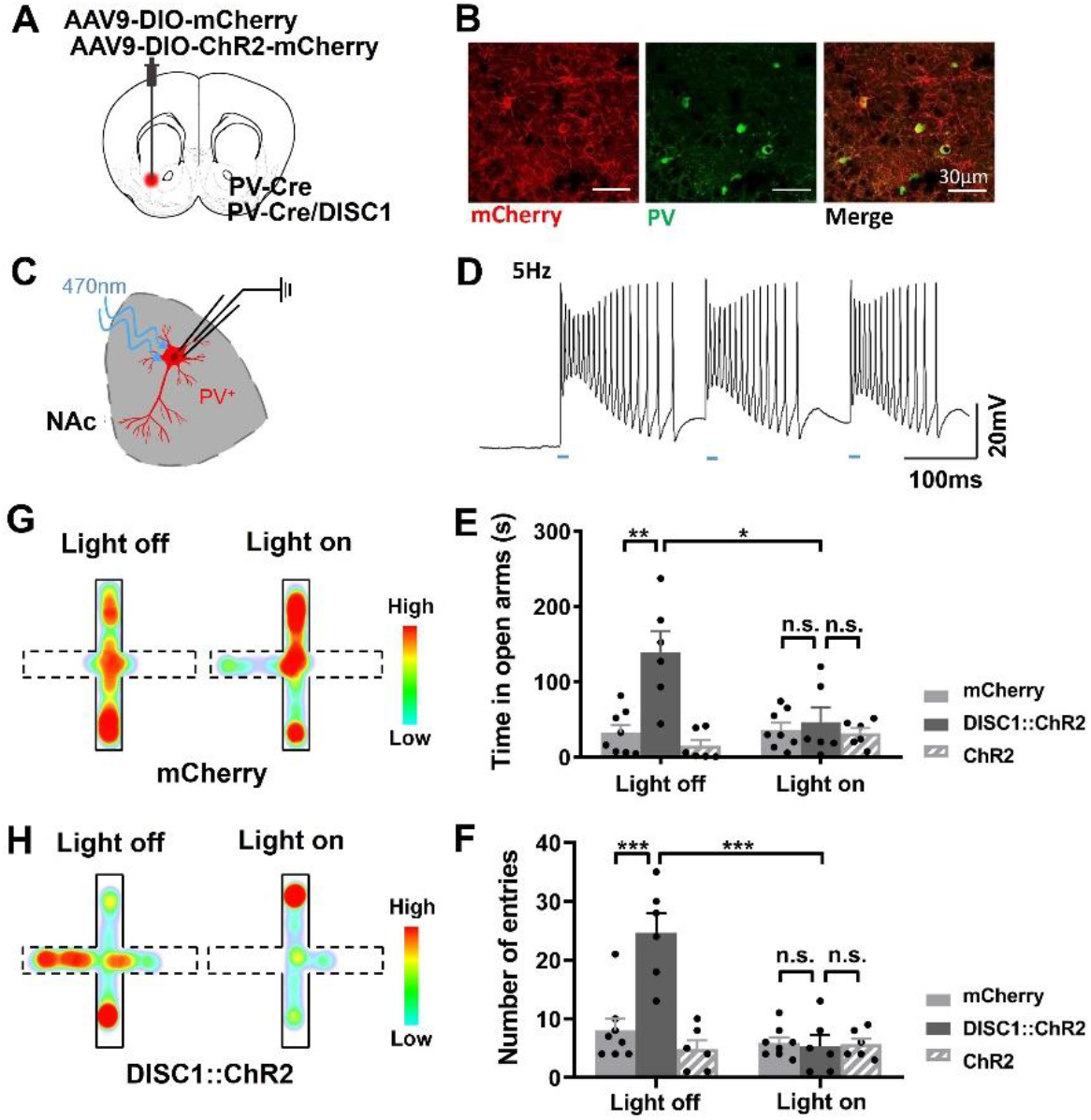
Optogenetic stimulation of NAc^PV^ neurons rescues risk avoidance deficiency in *DISC1-N^TM^* mice. A. Schematic showing the injection of virus. B. Neurons in the NAc infected with AAV-DIO-ChR2-mCherry (red) co-stained with PV neurons (green). C. Schematic showing patch clamp recording of a PV neuron in an acute brain slice. D. Schematic showing light-evoked action potential of NAc PV neurons. E,F. Time spent in the open arms (E) and number of entries to the open arms (F) before and after blue light (470 nm) stimulation during EPM test (top: \**P* =0.212, \*\**P* =0.0017, bottom: \*\*\**P* =0.0007 (left) and 0.0005 (right); n=8, 6 and 6 from left to right). G, H. Representative heatmap illustrating the time spent by a PV-Cre mouse in the open and closed arms in the EPM (G) and a PV-Cre/*DISC1-N^TM^* mouse (H) before and after blue light (470 nm) stimulation (dotted line represents the open arms).

Of the neurons in the NAc, 95% are medium spiny GABAergic neurons (MSNs). We tested whether activation of GABAergic neurons in the NAc could rescue abnormal risk-avoidance behavior in *DISC1-N^TM^* mice. By crossing Vgat-ChR2 mice with *DISC1-N* truncation transgenic mice, we labeled GABAergic neurons with ChR2 (Fig. S1 A, B). Following tamoxifen injection, *DISC1*/Vgat-ChR2 double transgenic mice spent more time in the open arms and had more open-arm entries than wild-type littermates (Fig. S1 C, D). Photostimulation of the NAc at 60 Hz led to a significant reduction in time spent in, and the number of entries to the open arms in *DISC1*/Vgat-ChR2 double transgenic mice, whereas compared to wild-type littermates, specific activation of NAc^GABA^ by light did not fully rescue the abnormal risk-avoidance behavior in *DISC1^TM^* mice (Fig. S1 C, D). This result further confirms that NAc^PV^ neurons play an important role in risk-avoidance behavior in *DISC1-N^TM^* mice during the EPM task.

### Inhibition of NAc PV neurons mimic the abnormal risk-avoidance behavior observed in *DISC1-N^TM^* mice

We have found that modulation of PV neurons in the NAc can rescue abnormal risk-avoidance behavior in *DISC1-N^TM^* mice. To assess the contribution of PV neurons in the NAc to the regulation of *DISC1-N^TM^* abnormal risk-avoidance behavior, we used designer receptors exclusively activated by a designer drug (DREADD), to inhibit the activity of NAc PV neurons in mice with normal DISC1 functions. PV-Cre mice were bilaterally infected with AAV9-DIO-hM4Di-mCherry (Figure 6A, B). After three weeks of recovery, mice were tested on elevated zero maze for avoidance behavior without CNO injection. Then, following a one-week interval, the same group of mice were given CNO and tested again on the elevated zero maze. We found that mice with hM4Di expressed in NAc^PV^ neurons had similar entry numbers compared to the first test but spent more time in the open segments (Figure 6C, D). Mean speed was similar before and after CNO administration (Figure 6E). This result suggests that this one intervention - inhibition of NAc PV neurons using hM4di - is able to mimic the abnormal risk-avoidance behavior of *DISC1-N^TM^* mice.

**Figure 6.**
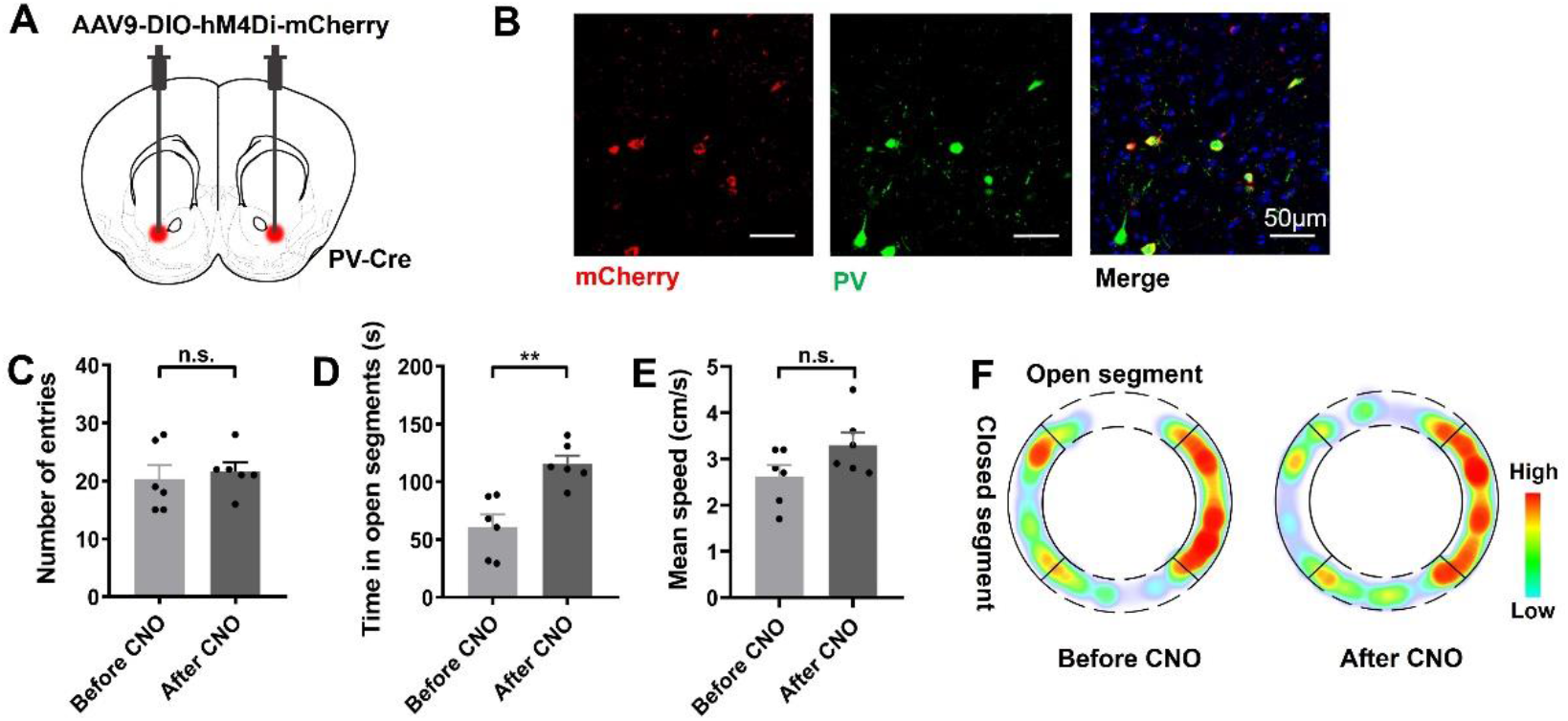
Inhibition of PV neurons activity in the NAc mimics *DISC1*-like abnormal risk-avoidance behavior. A. Schematic showing the injection of virus. B. Neurons in the NAc infected with AAV-DIO-hM4Di-mCherry (red) co-stained with PV neurons (green). C-E. Number of entries to the open segments (C), time spent in the open segments (D) and mean speed (E) before and after CNO administration during EOM test (\*\**P* =0.0017; n=6). F. Representative heatmap showing time spent in each location in EOM test before and after CNO administration.

## Discussion

In this study, we tested risk cognition behavior in *DISC1-N^TM^* mice and found a risk-avoidance impairment in the EPM test. This test is an approach-avoidance conflict task where anxious mice tend to avoid the open arms and favor the closed arms (‘safe’ zones) more so than non-anxious mice [32, 33]. Rodents have a natural aversion for open and elevated areas, as well as natural spontaneous exploratory behavior in novel environments. The EPM relies on the rodents’ preference for closed spaces (approach) to open spaces (avoidance) [34] and was directly used to determine the risk avoidance behavior [35]. Taking into account the impairments in social preference in *DISC1-N^TM^* mice, we predicted that these mice would spend more time in the open arms due to impaired normal cognition of the risks compared to their wild-type littermates. We also use an elevated zero maze to determine whether inhibition of NAc^PV^ neurons can mimic abnormal risk-avoidance behavior seen in *DISC1-N^TM^* mice. The elevated zero maze is a modification of the EPM that has the advantage of lacking ambiguity in scoring present in the EPM central area. When NAc^PV^ neurons in PV-Cre mice were inhibited, mice spent more time in the open segments but the number of entries to the open segments was not different from controls. The increased time spent in the open segments indicates an impairment in avoiding potential risks. This suggests that NAc^PV^ neurons are indispensable and are able to induce abnormal risk avoidance behavior of *DISC1-N^TM^* mice.

In order to find out which brain region is involved in the risk-avoidance behavior of *DISC1-N^TM^* mice after EPM tests, we measured neuronal activity in eight brain regions using c-Fos as an index. We found four brain regions where c-Fos was expressed significantly more than in WT mice: the NAc, BLA, PVT and the VTA were increased (in *DISC1-N^TM^* mice, compared with WT mice), suggesting that these regions may be involved in risk avoidance. In a structural investigation of mutant *DISC1* mice, it was found that the NAc expressed an abnormal dopamine type-2 receptor (D2R) profile and medium spiny neurons had reduced spine density on their dendrites [36]. In mutant *DISC1* mice, methamphetamine (METH)-induced dopamine (DA) release was significantly potentiated in NAc [37], and homeostasis of coincident signaling of DA and glutamate was altered within the NAc [38]. Considering these changes in the NAc due to the mutant *DISC1* gene, we decided upon the NAc as our target region to investigate the abnormal risk-avoidance behavior of *DISC1-N^TM^* mice.

Neurons with the capacity to discharge at high rates are called FS-neurons [39]. We found that FS-neurons in the NAc had reduced firing rates after tamoxifen administration and this correlated with a decrease in local theta power. In the NAc area, FS interneurons modulate principal neurons through its powerful and sustained feedforward inhibition [40, 41]. PV expressing interneurons, which constitute 1–2% of NAc neurons, have particular firing properties and are classified as FS-neurons [40, 42]. Other interneurons in the NAc that express somatostatin, neuropeptide Y, and neuronal nitric oxide synthase are classified as persistently low-threshold spiking (PLTS) neurons [42]. Although many studies have reported FS neurons in the NAc as PV neurons, PV interneurons in the NAc are not homogeneous. One study discovered that a portion of NAc FS-neurons also express cannabinoid receptor 1 (CB1) [43, 44]. These NAc^CB1^ neurons partly overlap with NAc^PV^ neurons (52%) [43]. Whether CB1-expressing FS-neurons have the same function as PV-expressing FS-neurons in the NAc, or whether CB1-expressing FS-neurons can been regarded as a subtype of NAc FS-neurons remains an important question. Considering that the striatal PV-expressing interneurons recorded thus far are FS-neurons [45, 46] and the functional properties of FS-neurons reported thus far have uniformity [44, 47, 48], we think PV can serve as a reliable marker for FS-neurons in the NAc. Using these guidances, we deduced that the FS-neurons we recorded in the NAc were PV neurons. We further explored the function of NAc^PV^ neurons by *in vitro* whole cell patch clamp and behavioral tests. We found the current-stimulated firing rate of NAc^PV^ neurons was lower in *DISC1-N^TM^* mice compared with control mice and light-evoked activation of PV neurons was able to rescue risk-avoidance impairment in *DISC1-N^TM^* mice. Both of these results are evidence for our deduction. Normally, NAc^PV^ neurons receive excitatory inputs from the same brain areas that project to NAc^MSN^ neurons (95% of total neurons in NAc) and form functional contacts with NAc^MSN^ neurons [47–49]. Whether NAc^PV^ neurons function in the same way in risk-avoidance impairment in *DISC1-N^TM^* mice has not been explored yet.

Although many different *DISC1* models have been generated, phenotypes have not always been consistent [6, 50], and as such, the influence of the *DISC1* gene on mental disorders remain elusive. As an important hub protein, DISC1 can interact with a number of synaptic or cytoskeletal molecules and modulate many cellular functions, such as synaptic plasticity [51] and neurogenesis [52, 53]. As such, different mutant models (e.g., *DISC1* mutation overexpressed model [8]) may influence DISC1 function to varying degrees and lead to different phenotypes. In this article, we used N-terminal fragment *DISC1* transgenic mice. Although we were focused on the cognitive impairments of the *DISC1-N^TM^* model, we cannot exclude the possibility that these *DISC1-N^TM^* mice also have abnormal anxiety states on the EPM. Anxiety, as an emotion, can act as a driver of decision making [54, 55]. More behavioral experiments are need to further investigate any causal relationship between risk avoidance impairment and abnormal anxiety states in *DISC1-N^TM^* mice.

In conclusion, we confirmed the cognitive impairment of *DISC1-N^TM^* mice in recognition of risks and found a lower firing rate in NAc^PV^ FS neurons both *in vivo* and *in vitro*. Reduced excitability of NAc^PV^ neurons may be responsible for the impairment in risk cognition and optogenetically increasing the activity of NAc^PV^ neurons can rescue risk-avoidance impairment in *DISC1-N^TM^* mice. These findings add to our understanding the neuronal circuits that relate to environment risk signals and their related evolutionary significance.

## Acknowledgements

This work was funded by the National Natural Science Foundation of China (31671116 J.T., 31761163005 J.T., 31800881 L.W. and 91132306 L.W.), the International Big Science Program Cultivating Project of CAS (172644KYS820170004 L.W.), the External Cooperation Program of the Chinese Academy of Sciences (172644KYSB20160057 J.T.), the Youth Innovation Promotion Association of the Chinese Academy of Sciences (2017413 P.W., Y6Y0021004 J.W.), the Guangdong Provincial Key S&T Program (2018B030336001 J.T.), Shenzhen Government Basic Research Grants (JCYJ20170413164535041 L.W.), Shenzhen Discipline Construction Project for Neurobiology DRCSM [2016]1379 (L.W.).

We thank Mr. Xu ZB and Mr. Liu BF for their help in transgenic mice husbandry and phenotyping. We are grateful to Ms. Li NN for the help in virus packaging.

## Author contributions

J.T. conceived of this study. X-Y.Z., B-F. W, Q.X., Y. Z., X. Z. and Y.L. performed experiments. J.T., X-Y.Z., P-F.W., W.H., and J.W. analyzed data. W-D.L. provided the *DISC1-N^TM^* mice and related materials. L-P. W. provided suggestions and comments on the manuscript. J.T., X-Y.Z. and B-F.W. wrote the manuscript.

---

**Suppl. Figure 1.**
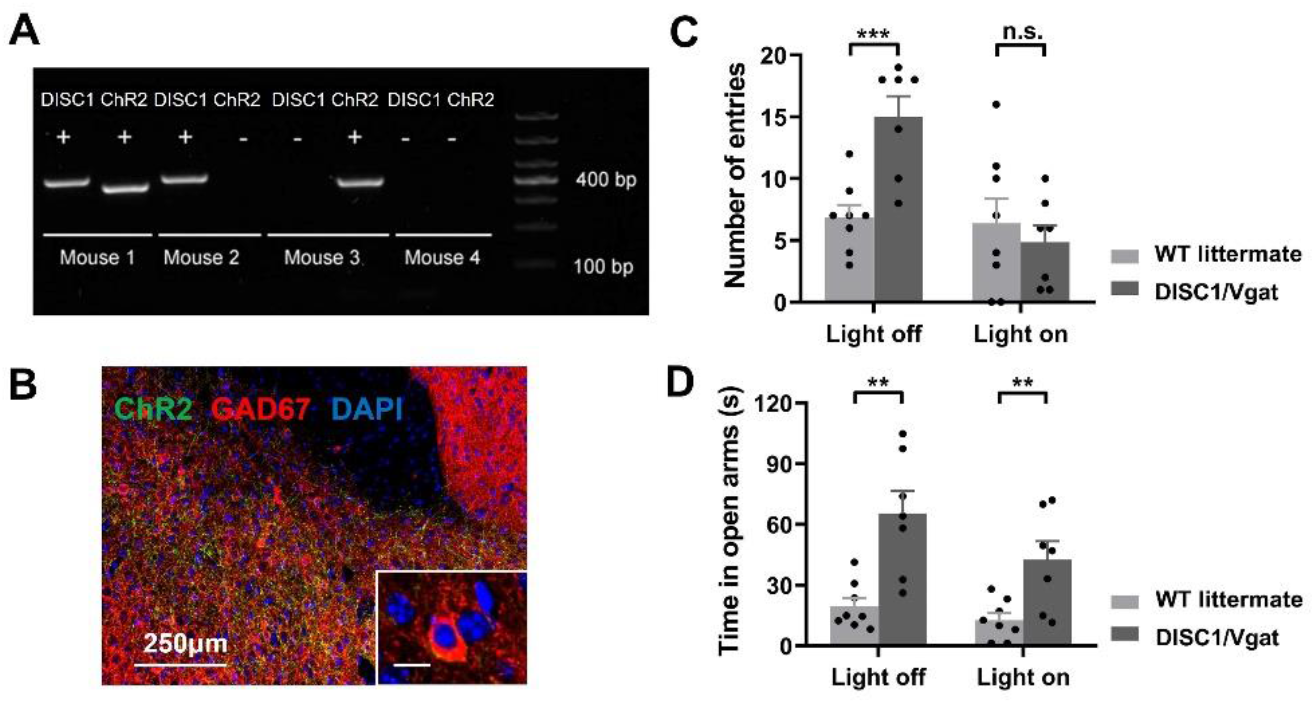
Optogenetic stimulation of NAc^GABA^ neurons did not fully rescue risk-avoidance imapairment in *DISC1-N^TM^* mice. A. Sample gel showing the successful generation of DISC1/Vgat-ChR2 double transgenic mice. B. Representative confocal image showing targeted ChR2 expression (green) co-stained with GABAergic neurons (red) in the NAc in these double transgenic mice. Scale bars= 250 μm, 10 μm (inset). C, D. Number of entries (C) and time spent in the open arms (D) before and after blue light (470 nm) stimulation during the EPM test (top: ***P =0.0009, bottom: **P =0.0014 (left) and 0.0066 (right); n=8 left, n=7 right).

